# Phosphatidylinositol 4,5-bisphosphate regulates cilium transition zone maturation in *Drosophila melanogaster*

**DOI:** 10.1101/315275

**Authors:** Alind Gupta, Lacramioara Fabian, Julie A. Brill

**Author notes:** To whom correspondence should be addressed: Julie A. Brill, Cell Biology Program, The Hospital for Sick Children, PGCRL Building, 15th Floor, 686 Bay Street, Room 15.9716, Toronto, Ontario M5G 0A4, Canada.

## Abstract

Cilia are cellular antennae that are essential for human development and physiology. A large number of genetic disorders linked to cilium dysfunction are associated with proteins that localize to the ciliary transition zone (TZ), a structure at the base of cilia that regulates trafficking in and out of the cilium. Despite substantial effort to identify TZ proteins and their roles in cilium assembly and function, processes underlying maturation of TZs are not well understood. Here, we report a role for the membrane lipid phosphatidylinositol 4,5-bisphosphate (PIP_2_) in TZ maturation in the *Drosophila melanogaster* male germline. We show that reduction of cellular PIP_2_ levels by ectopic expression of a phosphoinositide phosphatase or mutation of the type I phosphatidylinositol phosphate kinase Skittles induces formation of longer than normal TZs. These hyperelongated TZs exhibit functional defects, including loss of plasma membrane tethering. We also report that the *onion rings* (*onr*) allele of Drosophila *exo84* decouples TZ hyperelongation from loss of cilium-plasma membrane tethering. Our results reveal a requirement for PIP_2_ in supporting ciliogenesis by promoting proper TZ maturation.

**Brief summary statement:** The authors show that the membrane phospholipid PIP_2_, and the kinase that produces PIP_2_ called Skittles, are needed for normal ciliary transition zone morphology and function in the Drosophila male germline.

## Introduction

Cilia are sensory organelles that are important for signalling in response to extracellular cues, and for cellular and extracellular fluid motility [1, 2, 3, 4]. Consistent with their importance, defects in cilium formation (*i*.*e*. ciliogenesis) are associated with genetic disorders known as ciliopathies, which can display neurological, skeletal and fertility defects, in addition to other phenotypes [5, 6, 7, 8]. Many ciliopathies are associated with mutations in proteins that localize to the transition zone (TZ), the proximal-most region of the cilium that functions as a diffusion barrier and regulates the bidirectional transport of protein cargo at the cilium base [9, 10]. For example, the conserved TZ protein CEP290 is mutated in at least six different ciliopathies [11] and is important for cilium formation and function in humans [12, 13] and Drosophila [14]. Although the protein composition of TZs has been investigated in various studies [15], the process of TZ maturation, through which it is converted from an immature form to one competent at supporting cilium assembly, is relatively understudied.

Ciliogenesis begins with assembly of a nascent TZ at the tip of the basal body (BB) [9]. During TZ maturation, its structure and protein constituents change, allowing for establishment of a compartmentalized space, bounded by the ciliary membrane and the TZ, where assembly of the axoneme, a microtubule-based structure that forms the ciliary core, and signalling can occur. In Drosophila, nascent TZs first assemble on BBs during early G2 phase in primary spermatocytes [16]. This occurs concomitantly with anchoring of cilia to the plasma membrane (PM), microtubule remodelling within the TZ [17, 18], and establishment of a ciliary membrane that will persist through meiosis [16] (Figure 1A). TZ maturation has been described in Paramecium [19], *Caenorhabditis elegans* [20] and Drosophila [18], and is most readily observed by an increase in TZ length in the Drosophila male germline.

**Figure 1.**
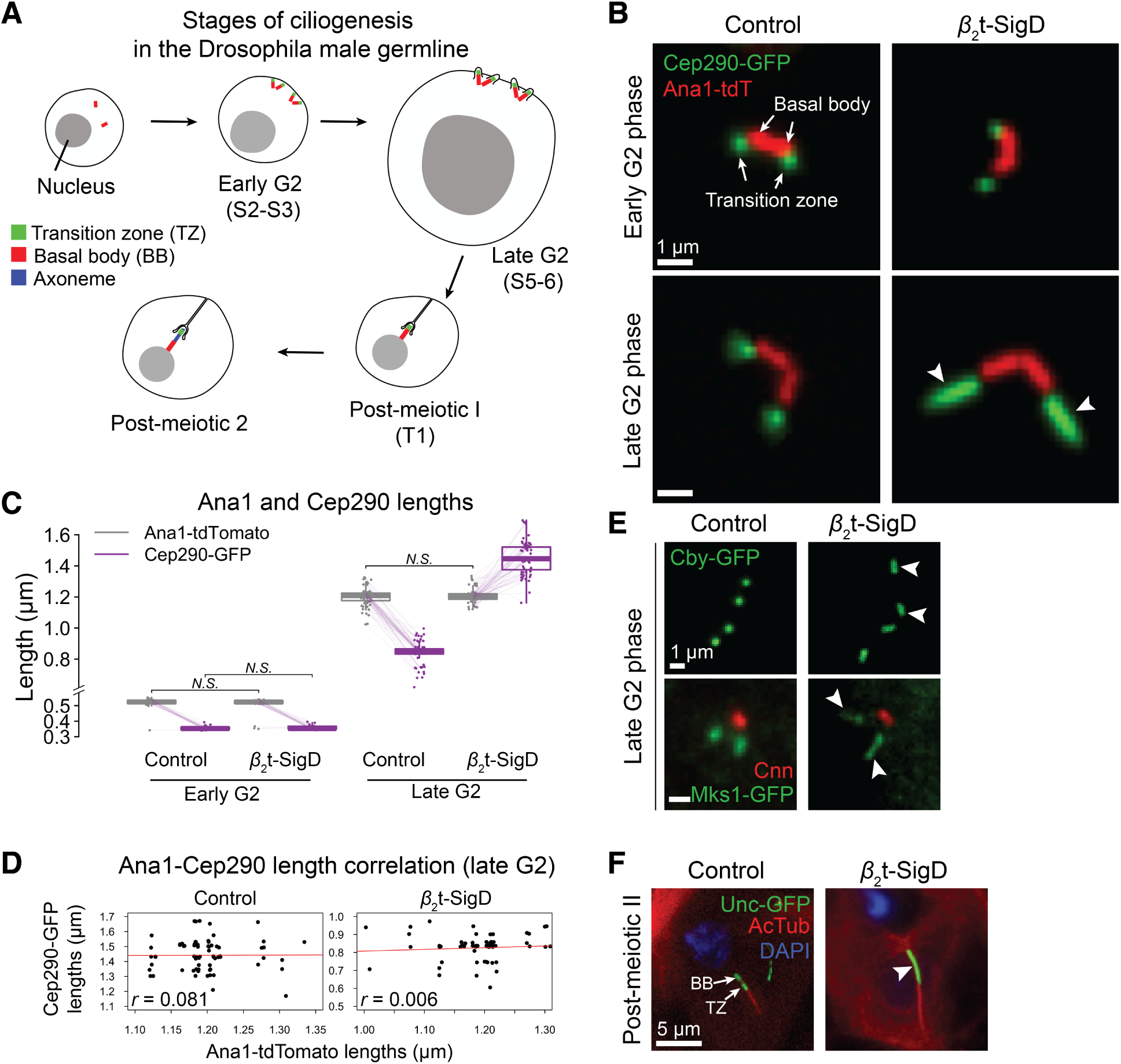
SigD expression induces transition zone hyperelongation. (A) Schematic diagram of the stages of ciliogenesis in the Drosophila male germline. Stages in parentheses correspond to those from [63]. (B) *β*_2_t-SigD expression induces Cep290 hyperelongation in cilia at late G2 phase (white arrowheads). (C) Quantification of paired Ana1-Cep290 lengths in early and late G2 phase in spermatocytes (*n* >30 and >65 respectively). (D) Lengths of Ana1-tdTomato versus Cep290-GFP in control and *β*_2_t-SigD cells at late G2 phase from (C) showing negligible correlation. Regression lines are plotted in red, and the Pearson correlation coefficient (*r*) is indicated on the bottom-left. (E) *β*_2_t-SigD expression induces hyperelongation of the TZ proteins Chibby (Cby) and Mks1 in late G2 phase (white arrowheads). (F) TZ hyperelongation persists through meiosis (white arrowhead) but does not affect axoneme outgrowth. Acetylated tubulin (AcTub) labels the axoneme.

We previously showed that the membrane lipid phosphatidylinositol 4,5-bisphosphate (PIP_2_) is essential for formation of the axoneme in the Drosophila male germline [21, 22]. PIP_2_, which is one of seven different phosphoinositides (PIPs) present in eukaryotes, localizes primarily to the PM, where it is required for vesicle trafficking, among other processes [23]. PIP_2_ has recently been linked to cilium function. Although the ciliary membrane contains very little PIP_2_ due to the action of the cilium resident PIP phosphatase INPP5E, the base of the cilium is enriched in PIP_2_ [24]. Inactivation of INPP5E causes a build up of intraciliary PIP_2_, which disrupts transport of Hedgehog signalling proteins in vertebrates [25, 26, 27] and ion channels involved in mechanotransduction in Drosophila [28]. In light of the current understanding of PIP_2_ as a modulator of cilium function, we sought to investigate the cause of defects we had observed in axoneme assembly in Drosophila male germ cells with reduced levels of PIP_2_ [21, 22].

## Results

### PIP_2_ is essential for transition zone maturation

To investigate how reduction of cellular PIP_2_ affects ciliogenesis in the Drosophila male germline, we used transgenic flies expressing the Salmonella PIP phosphatase SigD under control of spermatocyte-specific *β*_2_-tubulin promoter (hereafter *β*_2_t-SigD) [21]. To examine whether axoneme defects in *β*_2_t-SigD [21] were caused by aberrant TZ function, we examined localization of fluorescently-tagged versions of the core centriolar/BB protein Ana1 (CEP295 homolog) [29, 30] and the conserved TZ protein Cep290 [14] during early steps of cilium assembly. Cep290 distribution appeared similar in control and *β*_2_t-SigD in early G2 phase, when TZs are still immature. In contrast, Cep290-labelled TZs were significantly longer in *β*_2_t-SigD compared to controls by late G2 phase, following the period of TZ maturation (Figure 1B and 1C). Unlike Drosophila *cep290* mutants, which contain longer than normal BBs [14], Ana1 length was not affected in *β*_2_t-SigD, and we did not observe a strong correlation between Cep290 and Ana1 lengths (Figure 1D). Consistent with this result, the ultrastructure of BBs in *β*_2_t-SigD is normal, and localization of the centriolar marker GFP-PACT [31] is similar, in controls and *β*_2_t-SigD [21]. In contrast, TZ proteins Chibby (Cby) [32] and Mks1 [33, 34] exhibited hyperelongation in *β*_2_t-SigD (Figure 1E), indicating that this phenotype is not unique to Cep290. TZ hyperelongation was a highly penetrant phenotype (>70%) and showed high correlation (>0.95) within syncytial germ cell cysts, suggestive of a dosage-based response to a shared cellular factor, presumably SigD. Despite persistence of hyperelongated TZs through meiosis, axonemes were able to elongate normally in post-meiotic cells (Figure 1F). Nonetheless, the ultrastructure of these axonemes is frequently aberrant, either lacking nine-fold symmetry or containing triplet microtubules in addition to the usual doublets [21].

### The type I PIP kinase Skittles regulates TZ length

Although PIP_2_ is its major substrate in eukaryotic cells *in vivo* [35, 36, 37], SigD can de-phosphorylate multiple PIPs *in vitro* [38]. To address whether TZ hyperelongation observed in *β*_2_t-SigD represented a physiologically relevant phenotype due to decreased PIP_2_, we at-tempted to rescue this phenotype by co-expressing *β*_2_t-SigD with fluorescently-tagged Skittles (Sktl) under control of *β*_2_-tubulin promoter. We found that Sktl expression was able to suppress TZ hyperelongation in a cilium-autonomous manner (Figure 2A and 2B). Furthermore, the BB/TZ protein Unc-GFP [39, 21], exhibited TZ hyperelongation at a low penetrance in *sktl*^2.3^ mutant clones (Figure 2C), indicating that Sktl is important for TZ maturation.

**Figure 2.**
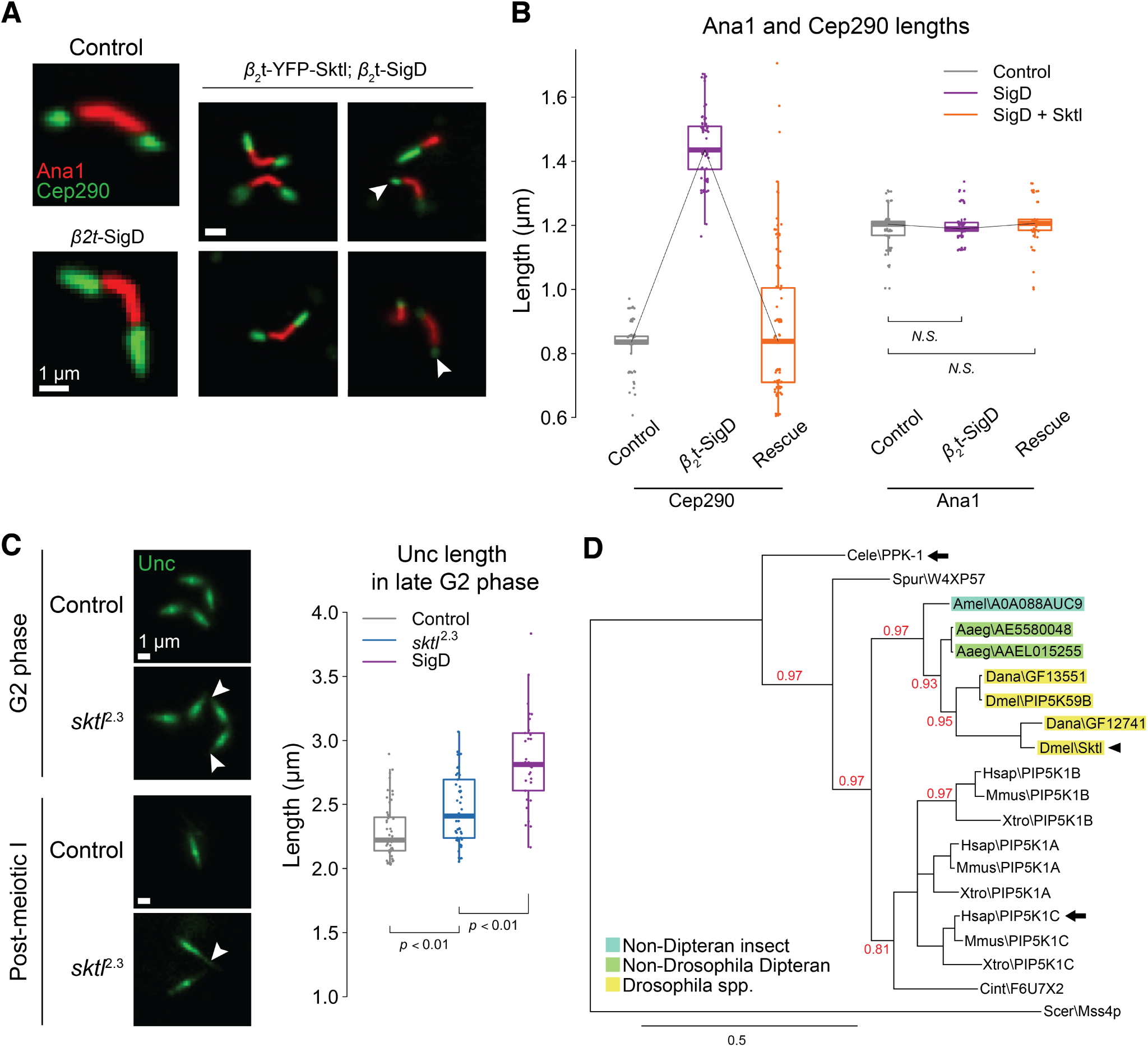
Sktl is important for transition zone maturation. (A) Expression of full-length Sktl can suppress *β*_2_t-SigD-induced TZ hyperelongation in a cilium-autonomous manner. Images were chosen to demonstrate varying levels of rescue of Cep290-GFP length in *β*_2_t-YFP-Sktl; *β*_2_t-SigD. White arrowheads mark rescued Cep290 distribution for comparison. (B) Quantification of Cep290 and Ana1 lengths from control, *β*_2_t-SigD from (A) and *β*_2_t-YFP-Sktl; *β*_2_t-SigD (*n* = 100). (C) *sktl*^2.3^ clones exhibit TZ hyperelongation (white arrowheads), as marked by Unc-GFP (left). Quantification of Unc-GFP lengths in control (*n* = 53), *sktl*^2.3^ (*n* = 31) and *β*_2_t-SigD (*n* = 51) spermatocytes at late G2 phase (right). (D) Phylogenetic tree of PIP5Ks showing evolutionary conservation of cilium-associated functions. Scale bar (bottom) represents expected amino acid substitutions per site. Branch support values are shown in red (a value of 1 indicates maximum support). Black arrows represent previous evidence of involvement in cilium-associated functions (from [40]). Black arrowhead indicates Sktl. Abbreviations: Cele (*Caenorhabditis elegans*), Spur (*Strongylocentrotus purpuratus*), Amel (*Apis mellifera*), Aaeg (*Aedes aegypti*), Dana (*Drosophila ananassae*), Dmel (*Drosophila melanogaster*), Hsap (*Homo sapiens*), Mmus (*Mus musculus*), Xtro (*Xenopus tropicalis*), Cint (*Ciona intestinalis*), Scer (*Saccharomyces cerevisiae*).

Vertebrate type I PIP kinase PIPKI*γ* has previously been shown to be important for cilium formation in cultured cells [40]. The two Drosophila PIPKIs, Sktl and PIP5K59B, arose from recent duplication of the ancestral PIPKI gene, and are not orthologous to specific vertebrate PIPKI isoforms (Figure 2D). Sktl has diverged more than its paralog PIP5K59B and seems to be functionally related to PIPKI*γ* and the *C. elegans* PPK-1 in having roles at cilia [41]. However, unlike the human PIPKI*γ*, which licenses TZ assembly by promoting CP110 removal from BBs [40], our results suggest that Sktl functions in regulating TZ length but not TZ assembly. Notably, neither inactivation nor overexpression of *cp110* affects cilium formation in Drosophila, and Cp110 is removed from BBs in early primary spermatocytes [42].

### Hyperelongated transition zones exhibit functional defects

We next sought to examine whether TZ hyperelongation due to SigD expression affected TZ function. Following meiosis in the Drosophila male germline, TZs detach from the BB and migrate along the growing axoneme, maintaining a ciliary compartment at the distal-most ~5*μ*m where tubulin is incorporated into the axoneme [14, 43]. As shown by Unc and Cep290 localization, TZs in *β*_2_t-SigD were frequently incapable of detaching from BBs and migrating along axonemes despite axoneme and cell elongation (Figures 1E, 3A and 3B). Indeed, the previously reported “comet-shaped” Unc-GFP localization in *β*_2_t-SigD [21] persists during cell elongation after meiosis (Figure 3A, bottommost panel) despite elongation of the axoneme (Figure 1E).

**Figure 3.**
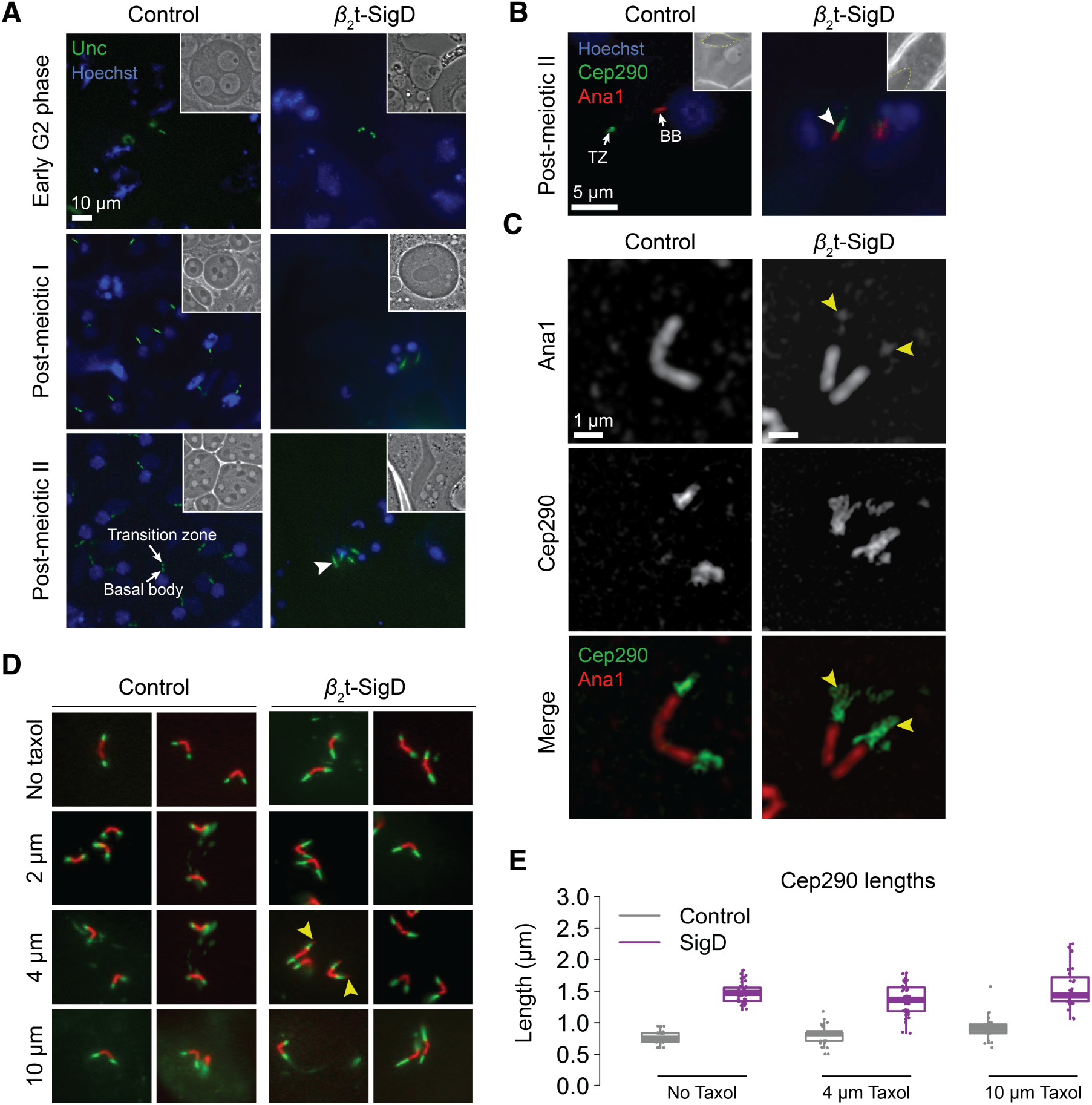
Hyperelongated transition zones display functional defects. (A) Unc-GFP is unable to detach from the basal body and migrate in spermatids expressing *β*_2_t-SigD (white arrowhead). Cell elongation in spermatids is concomitant with elon-gation of the mitochondrial derivative (dark organelles in the phase-contrast images). Insets show phase-contrast images corresponding to the region shown in the fluorescence images. (B) Cep290 is unable to detach and migrate from the basal body at the onset of axoneme assembly in *β*_2_t-SigD spermatids (white arrowhead). Insets show phase contrast images corresponding to the region shown in the fluorescence images, with the elongating mitochondrial derivative delineated by a yellow dashed line. (C) Structured illumination micrographs of *β*_2_t-SigD cells showing TZ-distal puncta of the centriolar protein Ana1 (yellow arrowheads). (D) Treatment of control and *β*_2_t-SigD cells with the microtubule stabilizing drug Taxol. Images demonstrate the variety in Cep290 distribution. Yellow arrowheads mark TZ-distal Ana1. (E) Quantification of Cep290 lengths in Taxol-treated control and *β*_2_t-SigD cells from (D) (*n* is between 30 and 40).

In Drosophila and humans, BBs consist of microtubule triplets [44, 45], whereas axonemes contain microtubule doublets due to obstruction of C-tubules at the TZ [18]. Consistent with a defect in this barrier and the presence of microtubule triplets in axonemes in *β*_2_t-SigD [21], a subset of cilia (<5%) in *β*_2_t-SigD contained puncta of Ana1 at the distal tips of TZs (Figure 3C). Treatment of germ cells with the microtubule-stabilizing drug Taxol increased the penetrance of this phenotype from <5% in untreated cells to >25% in cells treated with 4 *μ*M Taxol (arrowheads in Figure 3D) without significantly affecting Cep290 length (Figure 3E). Taxol-treated controls did not exhibit TZ-distal Ana1 puncta (*p* <0.01 at 5% penetrance). Fluorescently-tagged Asterless (CEP152 homolog), a pericentriolar protein [46, 47], did not localize to TZ-distal puncta in *β*_2_t-SigD (*p* <0.01) suggesting that these TZ-distal sites are not fully centriolar in protein composition. Taxol has been hypothesized to disrupt TZ maturation by inhibiting microtubule remodelling in the Drosophila male germline [17]. Similar to *β*_2_t-SigD, Taxol-treated male germ cells assemble extremely long axonemes that contain triplet microtubules [17], further supporting a functional relationship between PIP_2_ and microtubule reorganization in TZ maturation.

### The onion rings (onr) mutant decouples defects found in cells with reduced levels of PIP_2_

Male flies homozygous for the *onion rings* (*onr*) mutant of Drosophila *exo84* are sterile and exhibit defects in cell elongation and polarity similar to *β*_2_t-SigD [21]. Exo84 is a component of the octameric exocyst complex, which binds PIP_2_ and regulates membrane trafficking at the PM [48]. To investigate whether defects in TZ hyperelongation could be explained by defective Exo84 function, we examined TZs in *onr* mutants. Unlike *β*_2_t-SigD, *onr* did not display hyperelongated TZs (Figure 4A), suggesting that Exo84 is dispensable for TZ maturation.

**Figure 4.**
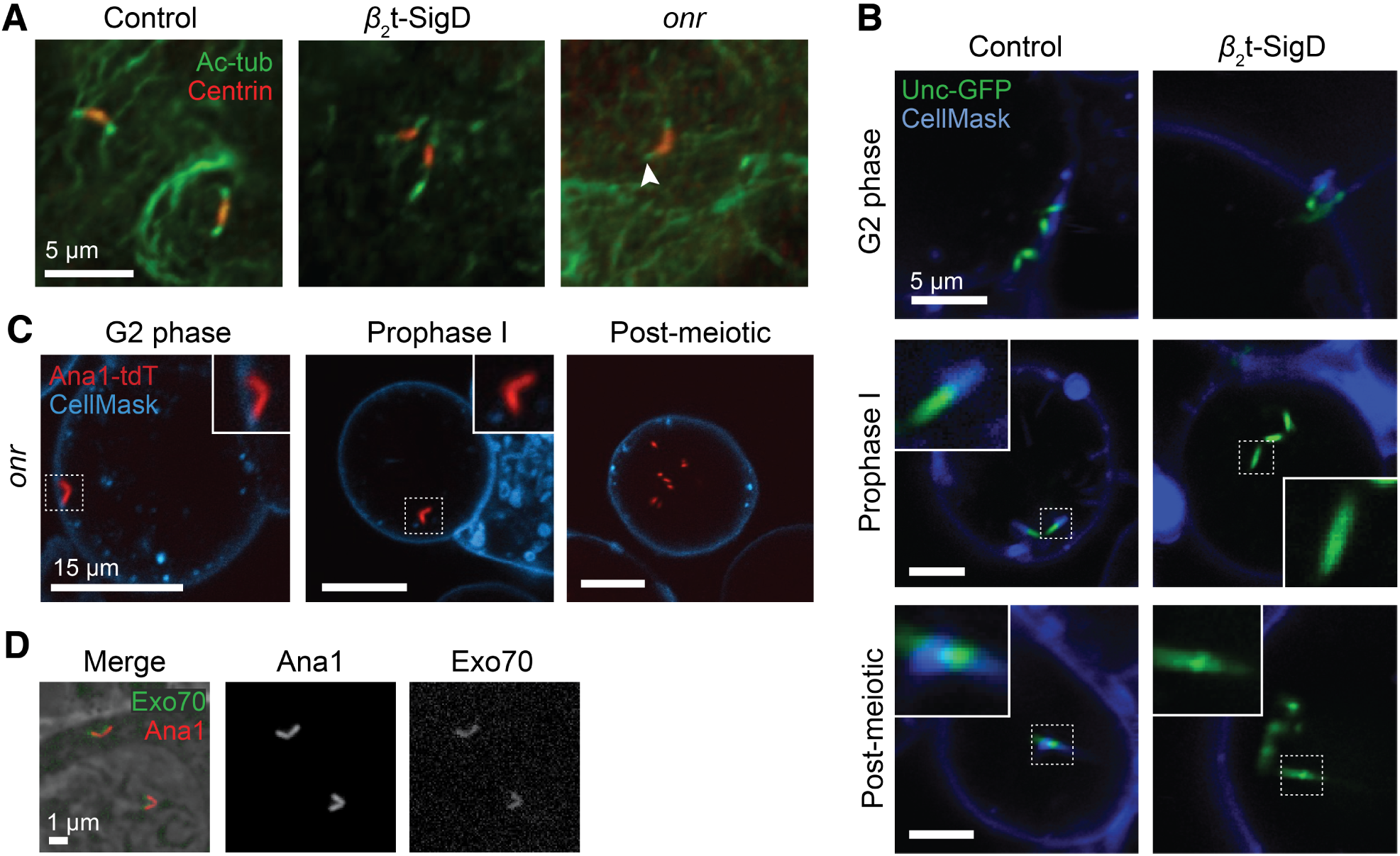
The onion rings (onr) allele of exo84 decouples TZ hyperlongation from loss of plasma membrane contacts. (A) *onr* mutant cells do not display hyperelongated acetylated tubulin signal at the cilium (white arrowhead). Acetylated tubulin marks the axoneme, which colocalizes with the TZ in spermatocytes [34]. (B) Cells expressing *β*_2_t-SigD fail to maintain cilium-PM tethering despite initially anchoring to the PM. The PM is marked with CellMask, a cell impermeable dye. (C) *onr* mutants do not maintain PM-cilium tethering. (D) GFP-tagged Exo70 localizes to BBs in spermatocytes.

Due to involvement of the exocyst in trafficking at the PM, we examined whether cilium-associated membranes were affected in *β*_2_t-SigD or *onr* mutants in a manner similar to *dilatory; cby* mutants [33]. Dilatory (Dila), a conserved TZ protein, cooperates with Cby to assemble TZs in the Drosophila male germline [33]. Whereas TZs in *β*_2_t-SigD and *onr* cells were able to dock at the PM initially, they were unable to maintain membrane connections, and were rendered cytoplasmic (Figure 4B and C), similar to TZs in *dila; cby* mutants. We found that fluorescently-tagged Exo70, a PIP_2_-binding exocyst subunit, localized to BBs (Figure 4D). Our results suggest that the exocyst, and Exo84 in particular, regulates cilium-PM associations, similar to PIP_2_, and that TZ hyperelongation and loss of cilium-PM association are genetically separable phenotypes.

## Discussion

The process of maturation of a TZ from a nascent form to a fully functional state, leading ultimately to axoneme assembly and ciliary signalling, requires orchestration of various proteins and cellular pathways [9, 15]. Our results indicate that normal execution of this process requires PIP_2_ and that depletion of PIP_2_ induces TZs to grow longer than normal. Similar to *β*_2_t-SigD, Drosophila *dila; cby* and *cby* mutants display hyperelongated TZs [32, 33], whereas *mks1* mutants have shorter TZs [34]. Because both Cby and Mks1 are hyperelongated in *β*_2_t-SigD cells, PIP_2_ regulates TZ length independently of an effect on Cby or Mks1 recruitment.

We also show that hyperelongated TZs are dysfunctional. Similar to *dila*; *cby* [33] and *cep290* [14] mutants, axonemes can assemble in *β*_2_t-SigD, albeit with aberrant ultrastructure [21], despite the lack of functional TZs or membrane association. The presence of TZ-distal Ana1 puncta in *β*_2_t-SigD cells, without the increase in BB length seen in *cep290* mutants lacking a functional TZ barrier, suggests that *β*_2_t-SigD expression selectively disrupts the ability of TZs to restrict C-tubules and Ana1 without abolishing the TZ barrier entirely. CEP295, the human Ana1 ortholog, regulates post-translational modification of centriolar microtubules [49], which may explain the presence of Ana1 along with supernumerary microtubules in *β*_2_t-SigD cells. Asterless (Asl), a pericentriolar protein important for centrosome formation and centriole duplication [46, 47], did not exhibit this TZ-distal localization, possibly due to differences dynamics of Ana1 and Asl loading onto centrioles [50, 51] or the more peripheral nature of Asl within the centriole [46].

The majority of PIP_2_ at the PM is produced by PIPKIs [23, 52]. In this study, we showed that mutation of the PIPKI Sktl induced hyperelongated TZs and that expression of Sktl could suppress TZ hyperelongation in *β*_2_t-SigD, with some cells showing cilium-autonomous suppression, suggesting Sktl might function *in situ* to regulate TZ length. In humans, *PIPKIγ* is linked to lethal congenital contractural syndrome type 3 (LCCS3), which has been suggested to represent a ciliopathy [40]. The recent discovery of a role for LCCS1-associated GLE1 protein in cilium function [53] corroborates this hypothesis. Our data support the idea that PIPKIs might represent ciliopathy-associated genes or genetic modifiers of disease.

Members of the exocyst complex such as Sec10 and Sec8 are important for cilium formation in cultured cell lines and zebrafish [54, 55, 56], but their precise roles in ciliogenesis are not well understood. The subunits Sec3 and Exo70 regulate exocyst targeting to the plasma membrane through a direct interaction with PIP_2_ [48, 57]. We previously showed that the *onr* allele of Drosophila *exo84* phenocopies defects in cell polarity and elongation observed in *β*_2_t-SigD [22]. Here, we show that the *onr* mutation phenocopies the loss of cilium-membrane contacts in *β*_2_t-SigD, similar to *dila; cby* mutants [33], but not TZ hyperelongation. Thus, TZ hyperelongation is not a prerequisite for the failure of cilium-PM association in male germ cells, and Exo84 uniquely regulates the latter process, potentially by supplying membrane required to maintain cilium-PM association. This result is supported by the Drosophila *cep290* mutant, which lacks a functional TZ but retains cilium-PM association [14]. Notably, *EXOC8*, which encodes the human Exo84, has been linked to the ciliopathy Joubert syndrome [58], and a similar process might underlie defects in humans with mutations in *EXOC8*.

## Methods

### Transgenic flies

Drosophila stocks were cultured on cornmeal molasses agar medium at 25°C and 50% humidity. Stocks expressing *β*_2_t::*Styp*\SigD (chromosome *3*) and *β*_2_t::YFP-Sktl (chromosome *2*) were described previously [21, 59]. GFP-Exo70 was cloned into the low-level expression vector *tv3* [59] and transgenic flies were generated using standard *P* element-mediated transformation. Ana1-tdTomato (chromosome *2*) and Cep290-GFP (chromosome *3*) were provided by T. Avidor-Reiss [14]. *Sp*/*CyO*; Unc-GFP was originally provided by M. Kernan [39]. Stocks expressing GFP-tagged Chibby and Mks1 were provided by B. Durand [32, 33]. The *onr* mutant was described previously [60]. Stocks for generating *sktl*^2.3^ clones were originally provided by A. Guichet [61]. Clones were induced by heat shock for two hours on days 3, 4 and 5 after egg laying. *w*^1118^ was used as the wild-type control.

### Antibodies

The following primary antibodies were used for immunofluorescence at the indicated concen-trations: chicken anti-GFP IgY (abcam), 1:1000; rat anti-RFP IgG (5F8, ChromoTek), 1:1000; rabbit anti-Centrin (C7736, Sigma-Aldrich), 1:500; mouse anti-acetylated *α*-tubulin 6-11-B (Sigma-Aldrich), 1:1000. Secondary antibodies were Alexa 488- and Alexa 568-conjugated anti-mouse, anti-rabbit and anti-chicken IgG (Molecular Probes) at 1:1000. DAPI at 1:1000 was used to stain for DNA.

### Fluorescence microscopy

For live imaging, testes were dissected in phosphate buffered saline (PBS). To stain for DNA, intact testes were incubated in PBS with Hoechst 33342 (1:5000) for 5 minutes. Testes were transferred to a polylysine-coated glass slide (Thermo Fisher Scientific) in a drop of PBS, ruptured using a syringe needle and squashed under a glass coverslip using Kimwipes. The edges of the coverslip were sealed with nail polish and the specimen was visualized using an epifluorescence microscope (Zeiss Axioplan 2) with an Axiocam CCD camera. Cells were examined live whenever possible to avoid artefacts from immunostaining.

For Taxol treatments, testes from larvae or pupae expressing Ana1-tdTomato; Cep290-GFP were dissected into Shields and Sang M3 medium (Sigma-Aldrich) supplemented with a predefined concentration of Taxol (Sigma-Aldrich) in DMSO and incubated overnight in a humidified sterile chamber in the dark at room temperature. These were then squashed in PBS and imaged live.

For CellMask staining, cells were spilled from testes in M3 medium onto a sterilized glass-bottom dish pre-treated with sterile polylysine solution to enable cells to adhere. CellMask Deep Red (Invitrogen) solution (20 *μ*g/mL) was added to the medium dropwise immediately before visualization under a confocal microscope.

For immunocytochemistry, testes were dissected in PBS, transferred to a polylysine-coated glass slide in a drop of PBS, ruptured with a needle, squashed and frozen in liquid nitrogen for 5 minutes. Slides were transferred to ice-cold methanol for 5-10 minutes for fixation. Samples were then permeabilized and blocked in PBS with 0.1% Triton-X and 0.3% bovine serum albumin, and incubated with primary antibodies overnight at 4°C, followed by three 5-minute washes with PBS, 1 hour incubation with secondary antibodies, and three 5-minute washes with PBS. Samples were mounted in Dako (Agilent) and imaged with a Zeiss Axioplan 2 epifluorescence microscope or a Nikon A1R scanning confocal microscope (SickKids imaging facility).

### Statistical methods

Statistical analysis and graphing was performed using R (version 3.4). A Gaussian jitter was applied when plotting results in Figures 1 and 2 for clearer visualization of trends, but raw data was used for all analyses. Statistical tests for “absence of phenotype” were computed using a binomial test under the assumption that the probability of the phenotype occuring was fixed. All *t*-tests were unpaired and two-sided with Welch’s correction for unequal variances. *n* represents the pooled number of samples (individual cilia) from multiple flies. A significance level of 0.01 was fixed *a priori* for all classical analyses. All raw data and code for analysis and plotting can be found online at http://www.github.com/alindgupta/germline-paper/.

### Phylogenetic analysis

Candidate orthologs of Skittles and PIP5K9B were queried from Inparanoid (version 8.0) and FlyBase (version FB2017_05). Poorly annotated protein sequences were confirmed to encode type I phosphatidylinositol phosphate kinases using reciprocal BLAST search. Phylogeny.fr (http://www.phylogeny.fr) [62] was used for phylogenetic reconstruction with T-Coffee for multiple alignment and MrBayes for tree construction. The output was converted to a vector image in Illustrator and colours were added for the purpose of illustration.

## Acknowledgements

We are grateful to Brian Ciruna for insightful discussions, to Bénédicte Durand, Tomer Avidor-Reiss, Antoine Guichet and Maurice Kernan for providing fly stocks, and to Bénédicte Durand and Bill Trimble for critical comments on the manuscript. This research was supported by a grant from the Canadian Institutes for Health and Research to J.A.B (MOP #130437). A.G. was supported by a University of Toronto Open Fellowship and an Ontario Graduate Scholarship.

## Author contributions

A.G. and J.A.B. conceived the project. A.G. performed all the experiments and analyses, and wrote the manuscript. L.F. generated GFP-Exo70 flies. J.A.B. edited the manuscript and supervised the project.

## List of Abbreviations

*β*_2_t-SigD: SigD driven by the male germline-specific *β*_2_-tubulin promoter
PIP_2_: phosphatidylinositol 4,5-bisphosphate
PIP: phosphoinositide, also known as phosphatidylinositol phosphate
TZ: transition zone
BB: basal body
PM: plasma membrane
*onr*: *onion rings* (an allele of *exo84*)

